# Ecological processes underlying the emergence of novel enzootic cycles—arboviruses in the neotropics as a case study

**DOI:** 10.1101/2020.04.24.057430

**Authors:** Sarah Guth, Kathryn Hanley, Benjamin M. Althouse, Mike Boots

## Abstract

Pathogens originating from wildlife (zoonoses) pose a significant public health burden, comprising the majority of emerging infectious diseases. Efforts to control and prevent zoonotic disease have traditionally focused on animal-to-human transmission, or “spillover”. However, in the modern era, increasing international mobility and commerce facilitate the spread of infected humans, non-human animals (hereafter animals), and their products worldwide, thereby increasing the risk that zoonoses will be introduced to new geographic areas. Imported zoonoses can potentially ‘spill back’ to infect local wildlife—a danger magnified by urbanization and other anthropogenic pressures that increase contacts between human and wildlife populations. In this way, humans can function as vectors, dispersing zoonoses from their ancestral enzootic systems to establish reservoirs elsewhere in novel animal host populations. Once established, these enzootic cycles are largely unassailable by standard control measures and have the potential to feed human epidemics. Understanding when and why translocated zoonoses establish novel enzootic cycles requires disentangling ecologically complex and stochastic interactions between the zoonosis, the human population, and the natural ecosystem. We address this challenge by delineating potential ecological mechanisms affecting each stage of enzootic establishment—wildlife exposure, enzootic infection, and persistence—applying existing ecological concepts from epidemiology, invasion biology, and population ecology. We ground our study in the neotropics, where four arthropod-borne viruses (arboviruses) of zoonotic origin—yellow fever, dengue, chikungunya, and Zika viruses—have separately been introduced into the human population. This paper is a step towards developing a framework for predicting and preventing novel enzootic cycles in the face of zoonotic translocations.

## Introduction

Humans have frequently enabled pathogens to overcome physical barriers to dispersal (1). The European conquest of the Americas brought Old World diseases to the New World, movement of troops during World War II propagated dengue viruses across the Asia-Pacific region (2), and air travel has provided an international transmission network for emerging infectious diseases (EIDs) such as the 2019 (ongoing) SARS-CoV-2 pandemic (3), the 2002-2003 SARS-CoV-1 outbreak (4) and pandemic influenza (5). Today, the majority of pathogens that infect humans are broadly distributed across geographic regions—globalized by human movement and population expansion, particularly during the past century (1). Animal pathogens have likewise spread globally through anthropogenic channels. The globalization of agriculture has expanded the geographic range of many livestock diseases with major economic repercussions, which continue to disproportionately affect the developing world (6). Domestic and wild animals translocated by humans have introduced their pathogens to new ecosystems, threatening biodiversity conservation—an anthropogenic impact termed “pathogen pollution” (7). In some cases, these invasive animal infections have maintained transmission post-emergence in local wildlife, establishing persistent reservoirs that subsequently reseed transmission and thwart control efforts in the original animal host population. Examples include African Swine Fever virus in Eastern Europe, where a novel enzootic cycle of the invasive livestock pathogen in wild boars has prevented disease eradication (8,9); and rabies virus in Africa, where human-mediated dispersal of domestic dogs established wild carnivore reservoirs that now contribute to rabies persistence in both wildlife and human communities (10).

Clearly, the global spread of zoonoses poses a unique and critical threat to human health. Novel enzootic cycles occur when zoonoses are introduced to new regions, infect local wildlife (spillback), and persist in local animal host populations (enzootic establishment). Figure 1 provides a diagram of these processes and Table 1 provides definitions of all terms in this paper). Now, more than ever, global conditions are ideal for the generation of novel enzootic cycles. In an increasingly connected world, international trade and travel provide pathways for pathogen introductions, while the recent surge in the emergence and re-emergence of animal pathogens has increased the number of zoonoses poised to exploit those pathways (11). Human population expansion into natural habitat is intensifying contact between humans and animals, creating more opportunities for imported zoonoses to spill back into naïve wildlife populations (12). The probability that these introduced infections persist in animal populations is increasing as human development pushes wildlife into crowded habitat patches and climate change alters transmission conditions (7).

**Figure 1.**
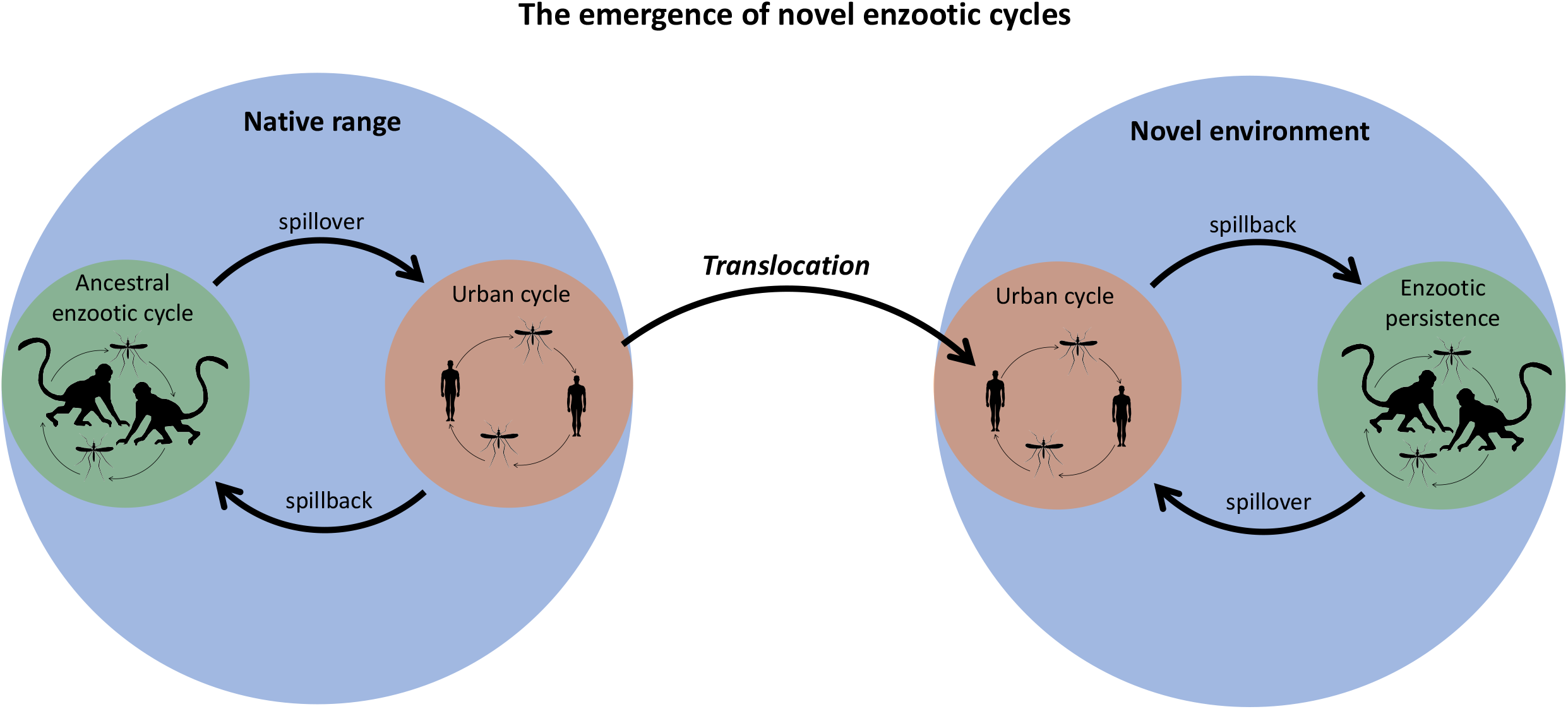
A diagram of the processes by which novel enzootic cycles emerge.

**Table 1.** Definitions of terms used in this paper.

| **Term** | **Definition** |
| --- | --- |
| Zoonosis | an animal pathogen that can also infect humans |
| Spillover | animal-to-human transmission |
| Spillback | human-to-animal transmission |
| Propagule pressure | the number, and temporal and spatial distribution of wildlife exposures to a translocated zoonosis |
| Permeability | the likelihood that humans and potential wildlife hosts, along with the translocated zoonosis, enter the boundary region at the human-wildlife interface |
| Realized niche | the set of hosts, vectors, and ecophysiological requirements that characterize existing transmission cycles of a translocated zoonosis |
| Fundamental niche | the set of hosts, vectors, and ecophysiological requirements that characterize the range of systems a zoonosis could theoretically invade if given the opportunity |
| Reservoir host | for a given zoonotic pathogen, the primary host species that is responsible for maintaining transmission |
| Enzootic (sylvatic) cycle | sustained and endemic transmission of a pathogen in a wildlife population |
| Enzootic (sylvatic) establishment | when a pathogen is introduced to a wildlife population and subsequently persists, attaining endemic status |
| Extrinsic incubation period (EIP) | the interval in the vector from initial infection until transmission is possible |

**Table 2.**
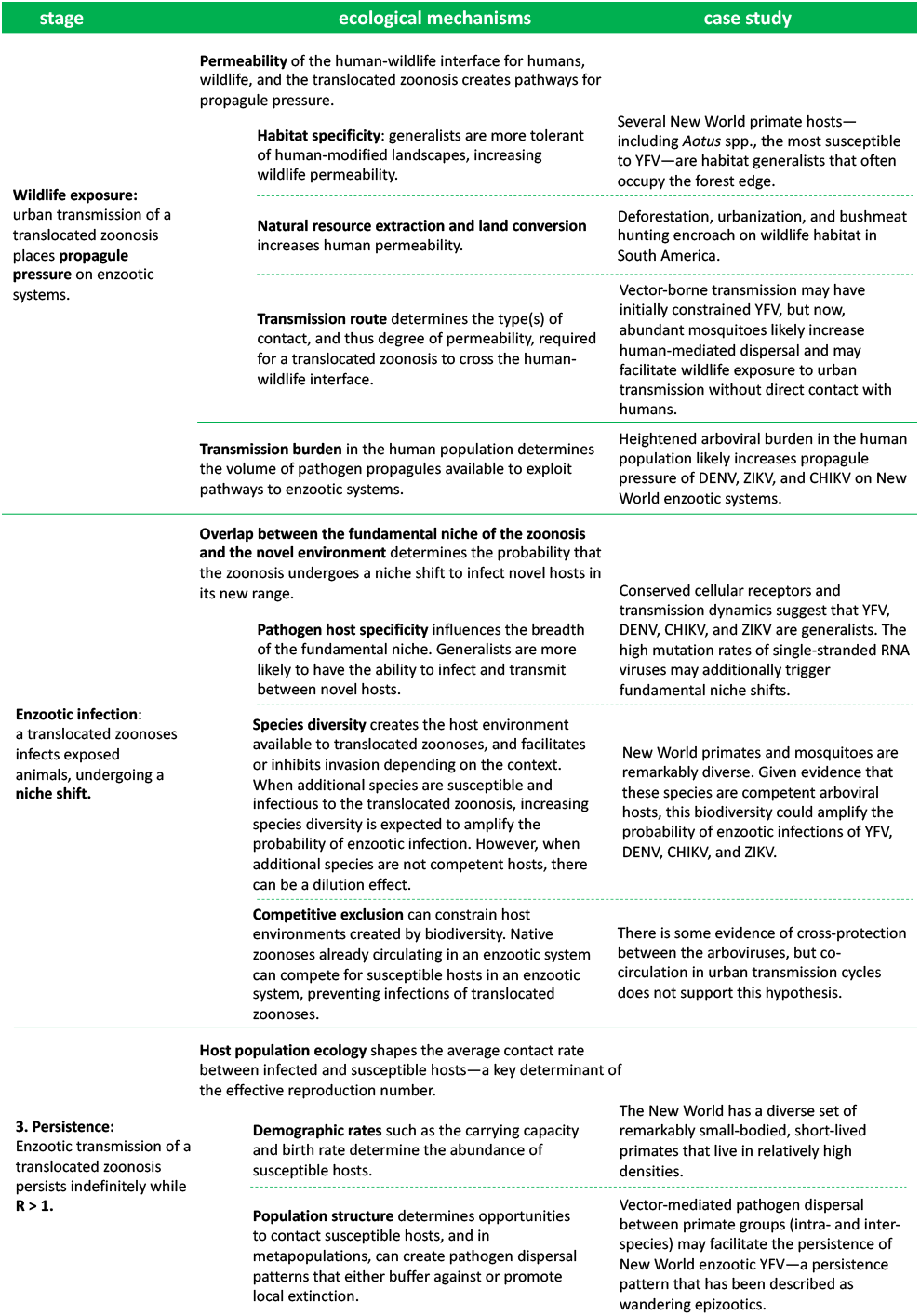
An outline of the ecological mechanisms affecting each stage of enzootic establishment, and their influence on the enzootic potential of translocated arboviruses in the neotropics.

In this era of globalization, zoonoses are increasingly being recognized as global threats. The emergence of SARS-CoV-2 in Wuhan, China has since affected 210 countries and territories, causing nearly 2 million cases and 125,000 deaths worldwide as of April 15^th^ 2020 (13). The pandemic has prompted an extraordinary global response—many countries have imposed nationwide lockdowns and closed their borders, nonessential international travel has largely been suspended (14), warring countries have declared cease-fire (15), and the World Health Organization (WHO) and United Nations have led international health and humanitarian organizations in mobilizing unprecedented funds for mitigating the spread and impact of the virus (16). Previously, the 2002-2003 SARS-CoV-1 epidemic prompted efforts to build infrastructure for global health security (17); and the WHO has declared recent outbreaks of Ebola, H1N1, and Zika as public health emergencies of international concern (PHEICs) (18). Nevertheless, this dialogue on the globalization of infectious disease continues to conflate zoonoses and human-specific pathogens, often overlooking what makes the spread of zoonoses so uniquely dangerous—the potential for enzootic reservoirs to establish in previously naïve regions. Some zoonoses such as Ebola virus and SARS-CoV-1 have remained within the human population after introductions to new regions. Conversely, introductions of yellow fever virus (YFV) in South America, *Yersinia pestis* (plague) in the Americas (19), rabies virus in parts of Africa, and West Nile virus in North America infected and persisted in local wildlife, inhibiting control and eradication efforts (10,20,21). The enzootic establishment of YFV in South America is a particularly noteworthy case because three New World introductions of three additional arboviruses—dengue (DENV), chikungunya (CHIKV), and Zika (ZIKV) viruses—have since followed. All three arboviruses now circulate in the same urban transmission cycle as YFV in the South American human population, raising the question: will their transmission remain within the human population, or will novel enzootic cycles emerge (22–25)? Identifying mechanisms that shape the outcome of zoonotic translocations is critical for developing strategies to mitigate the public health consequences of global transmission networks.

An emerging body of literature is beginning to discuss the risk that human-to-animal transmission will seed persistent enzootic reservoirs (22–25), but overall, disease emergence and spillover from wildlife continues to dominate the conversation on zoonotic transmission. Our understanding of enzootic establishment is limited by the difficulty in quantifying a process that is both highly stochastic and the product of interactions between multiple systems. In this review, we provide a conceptual framework to begin disentangling this ecological complexity, applying concepts from disease ecology, invasion biology, and population ecology. Cross-species pathogen emergence has previously been compared to species invasions (26) and population ecology used to understand post-introduction persistence (27) in the context of zoonotic spillover in human populations and host shifts within wildlife communities. Adapting this interdisciplinary theory on pathogen emergence to enzootic establishment, we review potential ecological mechanisms affecting the probability that translocated zoonoses emerge in novel enzootic cycles. We discuss the impact of each mechanism on the process of enzootic establishment: *(1)* local wildlife becomes exposed, *(2)* the zoonosis successfully infects the novel hosts, and *(3)* transmission persists indefinitely (28,29). We ground our discussion in the neotropics, where four arboviruses of zoonotic origin—YFV, DENV, CHIKV, and ZIKV—have separately been introduced into the human population. We additionally discuss the utility of modeling approaches, which we illustrate by building a simulation model for our neotropics case study. Our aim is to delineate the ecological processes that shape the outcome of zoonotic translocations as a first step towards developing a framework for predicting and preventing novel enzootic cycles, and therefore we do not discuss other key factors such as immunity, phylogeny, and evolution.

## The case study

YFV, DENV, CHIKV, and ZIKV all originated in sylvatic cycles involving non-human primates and primatophilic *Aedes* spp. mosquitoes in either Africa or Southeast Asia. As a result of human introductions, all four now circulate in human urban cycles in the Americas vectored by the anthropophilic mosquitoes *Ae. aegypti* and *Ae. albopictus* (30). YFV eventually spilled back to infect New World mosquitoes and primates, establishing a novel enzootic cycle that is broadly similar to the sylvatic transmission cycle in which it originated, albeit the taxonomic separation between Old World and New World hosts and vectors (31). To date, only YFV has successfully established sustained sylvatic transmission in New World primate and mosquito populations (31). However, given the similar histories of DENV, CHIKV, and ZIKV to that of YFV, there seems a high risk that neotropical mosquitoes and primates will also provide a suitable ecological niche for novel enzootic reservoirs of DENV, CHIKV, and ZIKV. DENV and ZIKV belong to the same genus, *Flavivirus,* as YFV, and CHIKV (family *Togaviridae,* genus *Alphavirus*) belongs to the same phylogenetic group as Mayaro virus (MAYV), an endemic South American zoonosis that circulates in primates and *Haemagogus* spp. (32). DENV additionally has spilled back at least once into a novel enzootic system—the virus established a persistent enzootic cycle in Africa after being introduced from Asia (33)—which offers a formidable warning of enzootic establishment in the American tropics.

ZIKV infections have recently been detected in New World primates, suggesting that the emergence of a persistent enzootic cycle could be imminent. Carcasses of free-living *Callithrix* spp. (marmosets) and *Sapajus* spp. (capuchins) were found to be infected with ZIKV strains of the ZIKV lineage currently circulating in the South American human population (34) and several monkeys have tested PCR-positive (35,36). Mosquito surveillance in Brazil additionally detected an amplicon of DENV in a pool of *Haemagogus leucocelaenus* (37). Sylvatic YFV now feeds recurring human epidemics in South America, allowing urban YFV transmission to continue despite vaccination campaigns. Thus, preventing the other three arboviruses, as well as any new introductions, from also establishing persistent enzootic reservoirs will be critical for forestalling further human morbidity and mortality from zoonotic transmission in the American tropics. However, the majority of the work on these imported arboviruses has focused on retrospective analysis of the conditions that enabled their introductions—particularly the global invasion of mosquito vectors *Ae. aegypti* and *Ae. albopictus—*and the transmission burden in the human population (38–41). Only a few recent papers have begun to discuss the threat of enzootic establishment (22–25,42). We add to this discussion by considering how each ecological mechanism identified in our review may affect the trajectory of DENV, CHIKV, and ZIKV in South America, using YFV as a frame of reference.

## Wildlife exposure

Once a translocated zoonosis has established in a new human system, there is an immediate risk that the pathogen spills back into local wildlife populations. The probability of spillback first depends on the rate at which wildlife is exposed, which can be captured by propagule pressure—a concept from invasion biology that represents the number, and temporal and spatial distribution of nonnative individuals introduced to a new system, and a key determinant of invasion success (43,44).

Propagule pressure hinges on introduction pathways between a source and recipient population. The propagule pressure of a translocated zoonosis on local wildlife will vary based on the availability of transmission pathways between the human (source) and wildlife (recipient) populations. Borrowing from landscape and movement ecology, Borremans et al. (45) identified permeability—the likelihood that source and recipient hosts, along with the pathogen, enter an ecosystem boundary region—as the ecological basis of pathways available for cross-species pathogen emergence across ecosystem boundaries. With respect to the human-wildlife boundary, permeability for translocated zoonoses will increase with wildlife tolerance of (or preference for) anthropogenically modified landscapes, and human communities’ proximity to the edge of a species’ habitat and incursions into natural habitat for resource extraction.

Host boundary permeability creates opportunities for contacts between infected source and recipient hosts, increasing propagule pressure on the recipient host population. Transmission route determines the type(s) of contact, and thus degree of permeability, required for the translocated zoonosis to cross the human-wildlife interface (45,46). Zoonotic introductions can occur via direct contacts such as bushmeat hunting or the wildlife trade— common in the developing world—or via indirect mechanisms such as environmental contamination—e.g., if infected bats leave saliva on forest fruits consumed by humans or shed excreta in the human environment (61,62,66). For directly transmitted zoonoses, propagule pressure on local enzootic systems will require sufficient permeability for humans and wildlife to come into close contact. Conversely, zoonoses that can survive outside their hosts will be less constrained by host boundary permeability. Domestic animals often intersect with both humans and wildlife and thus, have the potential to bridge transmission between the two populations. Vectors can likewise function as bridge hosts; as long as a competent vector is present, vector-borne zoonoses only require some degree of spatial and temporal overlap to transmit between humans and wildlife.

Transmission burden in the human population inhabiting the new region—a combination of time since introduction and the number of subsequent human cases—will determine the volume of pathogen propagules available to exploit transmission pathways to local wildlife (48). Consistent circulation and a high number of cases in the human population may result in more opportunities to spill back into enzootic transmission cycles, producing greater propagule pressure. The precise propagule pressure that led YFV to invade New World non-human primate populations is unknown, but phylogenetic analyses suggest multiple spillback introductions (49,50). Evidence that YFV reached its current widespread distribution in South America through long-distance, human-mediated dispersal implies that many spillback introductions occurred across a broad spatio-temporal landscape.

Vector-borne transmission may limit opportunity for spillback introductions to wildlife populations during periods of robust vector control efforts. Notably, in the 20^th^ century, *Ae. aegypti* eradication campaigns significantly reduced the YFV burden, to the point where health officials erroneously considered the arbovirus to be eradicated from the New World (21). Following relaxation of eradication efforts, *Ae aegypti* populations rebounded and *Ae. albopictus* invaded (38). These vectors increase arboviral boundary permeability, allowing zoonotic exchange to evade physical barriers (i.e., animals and humans do not need to interact directly or even occupy the same habitat for effective contacts to occur) (51). *Ae. albopictus* populations are often peridomestic, which may further bridge urban and sylvatic systems (42). It has been hypothesized that human-mediated movement of infected vectors played a significant role in the spread of enzootic YFV (50). Human-mediated vector dispersal and vector capacity to transmit between species without direct contacts have the potential to similarly facilitate enzootic invasion of DENV, CHIKV, and ZIKV.

Several New World primates are considered habitat generalists, with high permeability of human-modified landscapes, likely increasing opportunities for human-to-animal pathogen introductions. In particular, *Aotus* spp. (howler monkeys)—the New World primate most susceptible to YFV—often occupy the forest edge and have been hypothesized to bridge urban and enzootic transmission of YFV (52). At the same time, deforestation, urbanization, and bushmeat hunting are pushing humans into wildlife habitat (53,54).

Inconsistent circulation in the human population likely limited YFV propagule pressure on New World non-human primates. A virulent pathogen, YFV historically emerged intermittently in large, deadly outbreaks in human cities, relying on re-introductions along shipping routes (21). However, in recent decades, unprecedented population growth combined with climate change and the re-invasion of *Ae. aegypti* has fueled an increase in the frequency and magnitude of arboviral epidemics in the neotropics (55). Between 1980 and 2007, the number of reported DENV cases in the Americas increased 4.6-fold (56,57) and in 2019, exceeded 3 million, surpassing a previous record of 2.4 million (58); the 2013 introduction of CHIKV resulted in 2.9 million cases within the following three years (59), and the 2015-2016 ZIKV pandemic reached 48 countries and territories (60). Furthermore, unlike YFV, DENV, ZIKV, and CHIKV lack available, safe vaccines (61). This heightened arboviral burden has the potential to increase propagule pressure of DENV, ZIKV, and CHIKV on New World wildlife populations, accelerating the timeline between translocation and enzootic establishment.

## Enzootic infection

Not all wildlife exposures result in enzootic infections. To progress to the second step of enzootic establishment, the translocated zoonosis must be able to infect the exposed animals. Infecting novel host species in a novel environment can be described as a niche shift. The concept of an ecological niche has many nuanced definitions in ecology, but generally represents the set of abiotic and biotic conditions that allow a species to occupy a particular space within an ecosystem (62). A pathogen niche is defined by its hosts, vectors, ecophysiological requirements, and the many ways in which these parts interact (63,64). Like other species, a translocated zoonosis will have a realized niche—existing transmission cycles—and a fundamental niche—the range of systems the zoonosis could theoretically invade if given the opportunity (63). The probability that the zoonosis undergoes a niche shift to infect novel hosts in its new range depends on the degree of overlap between its fundamental niche and novel environment (62).

Host specificity influences the breadth of the pathogen’s fundamental niche. Generalists are defined by broad fundamental host ranges (65), which intuitively, will intersect with a wider range of enzootic systems, facilitating shifts to novel hosts (66–68). Alternatively, zoonoses can evolve to invade environments outside of their original fundamental niches (62). Some pathogen types inherently have higher potential for fundamental niche shifts than others. In particular, YFV, DENV, CHIKV, and ZIKV are all single-stranded RNA viruses—a group of pathogens previously shown to be the most likely to shift host species, predisposed to cross-species emergence by high mutation rates (28). Species diversity creates the host environment available to a translocated zoonosis. Disease ecologists have described a complex relationship between biodiversity and pathogen transmission within a given focal host species, where increasing diversity can have either an amplification or dilution effect on transmission, largely contingent on changes in community capacity to support infection (community competence) (69,70). Similarly, biodiversity in enzootic systems may either facilitate or inhibit the invasion of a translocated zoonosis depending on the abundance and distribution of competent hosts and vector species— susceptible to the zoonosis and infectious enough to transmit to the next susceptible individual. When increasing biodiversity adds competent host and vector species, we expect an amplified probability that wildlife exposures result in enzootic infections. However, adding low-competence species may cause a dilution effect, where wildlife exposures are increasingly “wasted” on hosts that cannot support infection. Additionally, competition among infectious agents can constrain host environments created by biodiversity. Native zoonoses already circulating in an enzootic system can compete for susceptible hosts, driving the competitive exclusion of a translocated zoonosis (71–73).

The “complete” host ranges of YFV, DENV, ZIKV, and CHIKV are not known, as it is practically infeasible to detect every enzootic infection. However, evidence of broad cell tropism and evolutionarily conserved viral entry strategies (74,75) suggest wide fundamental host ranges which are more likely to launch into novel systems (76). Phylogenetic analysis has demonstrated the relative ease with which YFV can shift between human and non-human primate host types, with multiple sylvatic strains circulating in the human population, but no evidence of major genetic adaptations between urban and enzootic transmission cycles (77).

A remarkably diverse assemblage of non-human primates and mosquitoes inhabit the neotropics. This biodiversity could amplify the probability of novel enzootic infections because New World primates and mosquitoes are competent hosts of a broad range of arboviruses (24,78). Previous meta-analyses suggest that the probability of cross-species emergence increases as the phylogenetic distance between novel and original host species decreases (i.e., as hosts become more closely related) (67,68,79–82). Thus, YFV, DENV, ZIKV, and CHIKV may be predisposed to shift to New World monkeys and arboreal mosquito vectors, which are phylogenetically related to the Old World monkeys and *Aedes* spp. that maintain sylvatic YFV, DENV, ZIKV, and CHIKV in Africa and Asia (31). That being said, New and Old World monkeys and mosquitoes could be divergent in critical immune factors and/or cell surface receptors involved in viral infection. Nevertheless, experimental infection has demonstrated that neotropical primates are competent hosts for ZIKV and DENV (83–86); neotropical *Haemagogus leucocelaenus* and *Aedes terrens* are competent vectors for CHIKV (25); and *Sabethes cyaneus* is a competent vector for ZIKV, though significantly less competent than *Ae. aegypti* (87). However, even if neotropical primatophilic mosquito species have limited capacity to vector DENV, ZIKV, or CHIKV, as humans encroach on forest habitat, anthropophilic vectors *Ae. aegypti* and *Ae. albopictus* could play an increasingly important role in sustaining enzootic cycles in peridomestic urban forests (42).

Evidence of cross-protective effects that could drive competitive exclusion of DENV, CHIKV, and ZIKV from New World enzootic systems is inconclusive. Speculation that hyperendemic DENV has conferred widespread cross-immunity against YFV in Asia—where YFV has remained absent despite the presence of suitable vectors and an entirely susceptible human population—has since been challenged (31,88). Additionally, there is no evidence of reciprocal cross-protective effects that would allow YFV to exclude DENV from New World enzootic systems (89). It has been hypothesized that ZIKV cross-protective immunity against DENV underlies the decline in DENV incidence following the first ZIKV outbreaks observed in Brazil (90) and Colombia (91); however, neutralization assays have not supported this hypothesis (92). A recent experimental infection study in mice suggests strong cross-protection of CHIKV against Mayaro (MAYV) virus—the endemic alphavirus that circulates in South American primates and *Haemagogus* spp. (93). However, there is no evidence of reciprocal cross-protective effects that would allow MAYV to exclude CHIKV from New World enzootic systems (89).

## Persistence

The outcome of spillback events depends on the potential for transmission between individuals in the novel animal host population. If animal transmission is limited, spillback might result in an isolated wildlife case, or alternatively, trigger an outbreak that threatens conservation efforts but eventually dies out (94,95). However, above a critical transmission threshold, the translocated zoonosis will persist indefinitely—the final step of successful enzootic establishment. In disease ecology and epidemiology, that critical transmission threshold is represented by the effective reproduction number (R)—the average number of secondary cases generated by a single infected individual in a population of susceptible and non-susceptible hosts. While R > 1, each infected individual will, on average, produce at least one secondary infection, allowing the pathogen to persist. The effective reproduction number is a function of pathogen attributes such as transmissibility and duration of infectiousness, as well as the average rate of contact between infected and susceptible individuals (and vectors if vector-borne) in a given population.

The average contact rate between infected and susceptible hosts is largely shaped by population ecology (96). Dynamic demographic rates and structuring of host populations determine the abundance, distribution, and movement of hosts available to translocated zoonoses in enzootic systems. Wildlife populations with high carrying capacities will have larger baseline populations of susceptible hosts. High birth rates replenish the supply of susceptible hosts, inhibiting mortality and conferred immunity from depleting the susceptible population and driving the effective reproduction below the threshold of persistence (97). Spatial structuring of susceptible hosts can either limit or enhance the potential for pathogen persistence. Pathogen dispersal between patches can promote persistence across a metapopulation by buffering against local depletion of susceptible individuals. On the other hand, spatial structure can limit contacts between patches, lowering the effective reproduction number and driving an epidemic to extinction (96,98).

While some New World monkeys are large species with low birth rates, one of the clade’s defining features is its diverse set of remarkably small-bodied, short-lived primates. For example, the pygmy marmoset (*Callithrix pygmaea*) is a small-bodied New World primate with bimodal annual birth peaks, high twin birth rates, and a natural lifespan of about 10 years (99). These smaller-bodied primates also tend to require smaller home ranges and live in higher densities, allowing for greater sympatric species richness. A meta-analysis of New World primate assemblages found that on average, forest sites contained six sympatric species, but this number could reach as high as 14 species, peaking near the equator (100). New World enzootic mosquito species occupy the canopy, preying primarily on primates. These vectors do not appear to demonstrate strong host preferences and thus, likely bridge transmission between sympatric groups of primate species. This spatial structure—vector-mediated pathogen dispersal between primate groups (intra- and inter-species)—may have facilitated the enzootic establishment of YFV. It has been hypothesized that enzootic YFV persists within primate metapopulations occupying continuous forest in wandering epizootics, in which transmission continually shifts between subpopulations. We suspect that the demographic rates and structuring of primate host and mosquito vector populations in the New World would, as with YFV, facilitate the persistence of DENV, CHIKV, and ZIKV should they spill back successfully.

## The ecology of enzootic establishment in the Anthropocene

Anthropogenic impacts are affecting ecological processes at every stage of enzootic establishment. Wildlife exposure to urban transmission of translocated zoonoses will likely increase as humans continue to encroach on wildlife habitats (2,12,22,23,101). Land conversion and extraction of natural resources drive humans, mosquito vectors, and primates to coincide in human-modified landscapes, potentially increasing the boundary permeability of the translocated arboviruses. Climate change will alter the epidemiology of zoonoses, particularly vector-borne zoonoses, substantially reducing transmission at temperature extremes, but increasing transmission in moderate warming scenarios (102). Rapid population growth and the global expansion of mosquito vectors contribute to heightened zoonotic transmission burden in the human population, which subsequently places greater propagule pressure on enzootic systems. Species composition is changing in many ecosystems, which can facilitate or inhibit pathogen invasions depending on whether lost species are competent hosts (70). However, habitat specialists are highly sensitive to anthropogenic impacts, whereas habitat generalists—which often bridge transmission between urban and enzootic systems—often persist (103). By increasing the concentration of competent hosts, anthropogenic change can have an amplification effect on enzootic infection. Habitat loss is additionally pushing animals into dense populations ripe for disease persistence (7).

Recent trends in enzootic YFV reflect the changing ecology of enzootic establishment in the Anthropocene. The past decade has seen a surge in the frequency and magnitude of YFV epizootics in South America. The discovery of co-circulating sylvatic transmission cycles during the 2016-2017 YFV epidemic in Brazil implies multiple spillback introductions on a local scale across a short timespan (77). Additionally, enzootic YFV has invaded previously non-endemic regions, significantly expanding its range in the New World.

## A case for modeling

It is challenging to approach the ecological complexity of enzootic establishment through field and experimental studies alone. Mathematical models can however be a useful tool that allows us to integrate existing data and ecological theory to elucidate system dynamics, particularly when data are sparse, as is often the case with enzootic systems (22,104,105). Many modeling approaches can be applied to a range of questions that allow us to better understand the risk that translocated zoonoses will emerge in novel enzootic cycles. For example, species distribution models, or ecological niche models—statistical approaches that leverage associations between presence-absence information and environmental variables to infer habitat suitability—can be used to identify at-risk systems where high permeability of humans, vectors, and translocated zoonoses create ideal conditions for enzootic establishment (46,106–108). Metapopulation modeling is often an important tool to demonstrate conditions of pathogen persistence in wildlife populations given that they are typically fragmented (96). Next-generation matrix methods are a useful tool to quantify the effective reproduction number of translocated zoonoses in novel enzootic systems to understand the probability of successful invasion (109). Explicit simulations that capture significant amounts of the complexity of systems have been effectively used to compare the impact of interventions in the human and wildlife population (110). We would also argue that simple models can be very useful in helping us understand the system-specific dynamics that influence the ecological processes underlying enzootic establishment. For example, Althouse et al. (22) developed the only previous model of vector-borne transmission in non-human primate hosts (22,104,105), and showed that whether ZIKV persists enzootically in South America is highly dependent on primate birth rates and mosquito population sizes. Based on model outcomes for the range of parameter values estimated to apply to the neotropics, they concluded that ZIKV has a high potential for enzootic establishment in the New World.

The modeling approach of Althouse et al. (22) can be applied to all four translocated arboviruses circulating in the neotropics. In particular, we can examine how key differences between the different viruses may impact their relative risk of enzootic establishment. Specifically, the extrinsic incubation period (EIP)—the delay from initial infection until transmission is possible—varies between YFV, DENV, CHIKV, and ZIKV (111–114). EIP helps determine the length of the mosquito infectious period, defining onward transmission (115) and therefore has the potential to impact the risk of spillback. However, a model is needed to explore whether these differences are likely to be relevant, particularly in the context of the long lifespans of New World sylvatic mosquitoes in South American enzootic systems. To examine this question, we built on the model of Althouse et al. (22), and simulated the introduction of a single infected primate within a multi-host metapopulation model to explore how a spillback event might play out for all four translocated arboviruses given New World mosquito lifespans (*see the Supplementary Information for a more detailed description of the model methods).* Except for the primate birth rate (which we hold constant), we reproduce their range of parameter values, additionally varying the length of the EIP to reflect differences between the arboviruses, and the mosquito lifespan to reflect differences between species and environmental conditions. Given that EIP and mosquito lifespan are highly variable across conditions, the values selected for our simulations were not meant to perfectly describe the four arboviruses or particular mosquito species; instead, we aimed to capture the range of values that have been observed across arboviruses and environmental conditions to understand general trends in the effect of EIP and lifespan on the probability of enzootic establishment. For EIP, we selected values between 2 (reflecting the lower bound for CHIKV (111)) and 10 days (approximating the longer EIP of ZIKV (87)). For mosquito lifespan, we use 7 days—taken from the previous model of enzootic vector-borne transmission in primates (22)—as our starting value and selected two additional values at 7 day intervals: 14 days approximately represents mean values from mark-release-recapture studies in South America for *Ae. albopictus* in field conditions (116), and 21 days approximately represents conservative mean values for *Haemogogus* spp. and *Sabethes* spp. at warmer temperatures (117). For each set of values, we ran 50 simulations, calculating the probability of persistence as the proportion of simulations in which infected primates remained at the end of the three-year period.

At shorter mosquito lifespans, EIP length affected the probability of persistence; increasing EIP to 10 days, as we might observe with ZIKV, decreased the probability of persistence, whereas shortening EIP to 2 days, as we might observe with CHIKV or at warm temperatures, increased the probability of persistence (Figure 2). However, extending the mosquito lifespan negated this effect; when the mosquito lifespan reflects New World sylvatic species and warm temperatures at 21 days, there is a relatively high probability of sylvatic establishment across all EIP values. Long-lived sylvatic mosquitoes are hypothesized to play a substantial role in YFV maintenance in the neotropics (118), which may be part of the reason why YFV has established and maintained enzootic transmission despite an EIP of 7 days. These results also suggest that despite differences in EIP, there may be a particularly high risk of enzootic establishment for DENV, CHIKV, and ZIKV in the New World due to naturally longer-lived sylvatic species and warmer temperatures, which both shorten EIP and extend mosquito lifespans. This work is important in highlighting the need for surveillance efforts to be equally vigilant of DENV, CHIKV, and ZIKV spillback in the New World. The results give a good example of how models can be useful in our understanding of complex ecological interactions. In this case we may have expected that difference in EIP may have been important in determining relative risk of enzootic establishment, however, due to longer mosquito lifespans, our model suggests that these differences are unlikely to be relevant.

**Figure 2.**
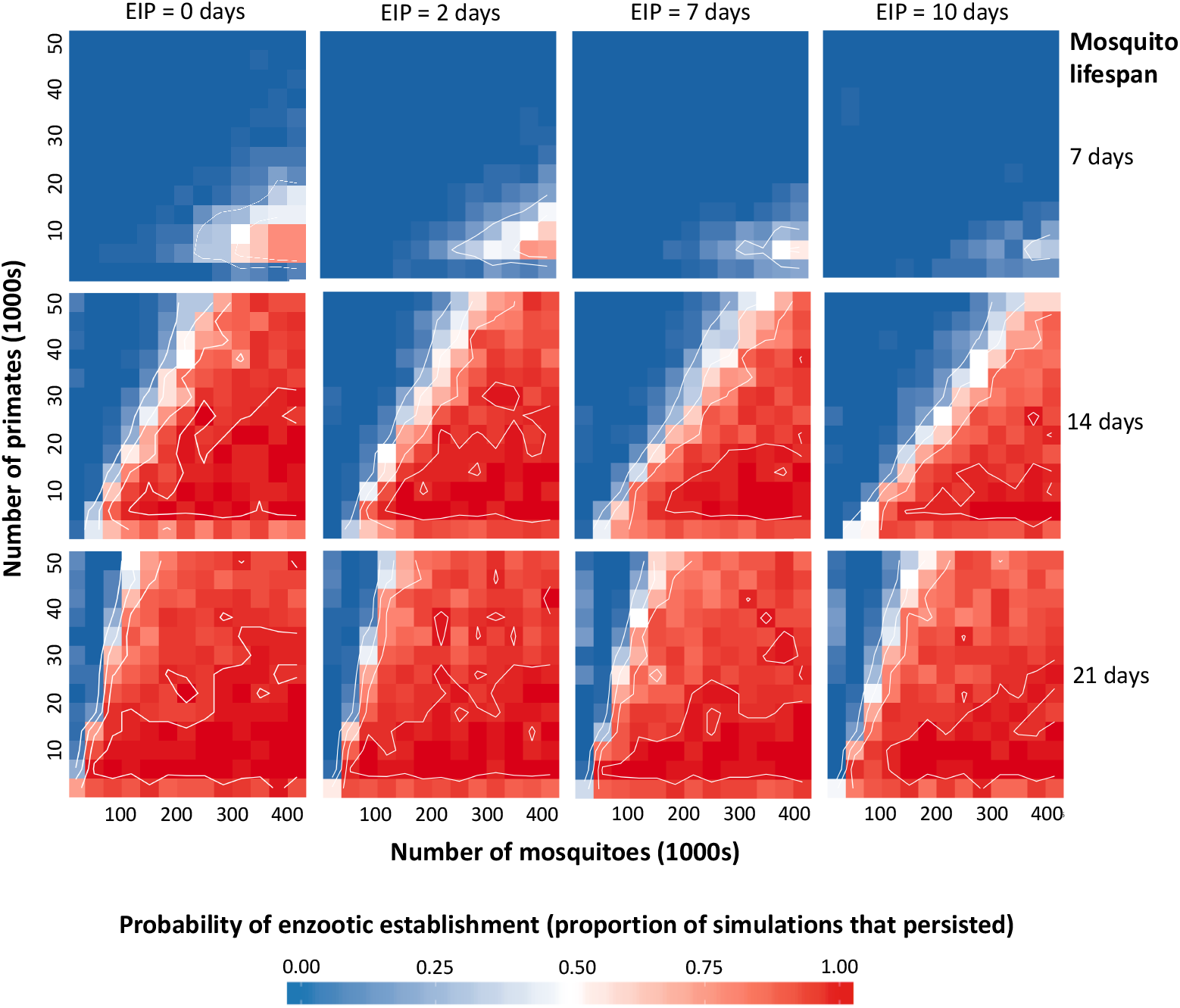
Model results predicting the probability of establishment across mosquito birthrates and EIPs. Mosquito birthrate is equivalent to 1/lifespan and increases from the top to bottom panels, while EIP increases left to right. Within each panel, the total population size of mosquitoes (in two populations) and primates (in two populations) changes horizontally and vertically, respectively. For each parameter set, we simulated the introduction of a single infected primate and subsequent transmission for a three-year period. Blue indicates no simulations establishing, whereas red indicates all simulations establishing. Contour lines show 0.25, 0.5, 0.75, and 0.95 probability of establishment.

## Discussion

The goal of this study is to stimulate research on the emergence of novel enzootic cycles and begin to disentangle the underlying ecology complexity. We have argued here that the establishment of novel enzootic cycles is a pressing threat with the capacity to dramatically alter disease dynamics. The International Task Force for Disease Eradication identifies the existence of an animal reservoir as a barrier to eradicating a disease because enzootic transmission often feeds human epidemics (119). In some cases, enzootic cycles have even contributed to the evolution of pandemic pathogens; for example, pigs have functioned as “mixing vessels” for the evolution of pandemic swine influenza (120). We delineated potential ecological mechanisms at each stage of enzootic establishment, grounding our discussion in the neotropics, where the danger of enzootic establishment is evident in the history of YFV and an ongoing threat given the endemic circulation of DENV, CHIKV, and ZIKV in the human population. Enzootic YFV, which has triggered devastating human epidemics across the neotropics, has expanded its geographic range since its initial establishment (121). There is a real danger that DENV, CHIKV, and ZIKV will also establish persistent enzootic reservoirs in the New World, similarly inhibiting efforts to prevent future human epidemics. Given that enzootic cycles are nearly impossible to control or eradicate, avoiding enzootic establishment will be critical to mitigate the current arbovirus public health emergency in the New World. Moreover, enzootic cycles can thwart efforts to eradicate pathogens in the human population, as has occurred with the carriage of Guinea worm by dogs (122). In particular, the recent discovery of natural ZIKV (34–36) infection in nonhuman primates in Brazil calls for renewed urgency to understand the potential for enzootic persistence.

Spillback events in which humans introduce pathogens into wildlife populations may become common occurrences, as urbanization and anthropogenic pressure on wildlife populations increase opportunities for human-wildlife contact. As a result, understanding the risk that arboviruses will persist in sylvatic cycles after spillback events should be established as a research priority, particularly in the era of SARS-CoV-2. There is significant concern that SARS-CoV-2 could spill back into susceptible wildlife within its expanded geographic range and establish novel enzootic reservoirs, becoming endemic outside of China. The high burden and global distribution of human SARS-CoV-2 transmission places propagule pressure on a wide range of enzootic systems. Bats—the putative origin of SARS-CoV-2 (123)—are the second most diverse mammalian group, inhabit every continent except Antarctica, and harbor a large diversity of coronaviruses (124). Many bat species migrate long distances (>1,000 km) (125), increasing their probability of exposure to SARS-CoV-2 and capacity to subsequently spread the virus. Although previous work has demonstrated that human-adapted CoV often cannot infect bat cells (126), the cell-surface receptor implicated in SARS-CoV-2 invasion of host cells—angiotensin converting enzyme 2 (ACE2)—has been found to be highly diverse across bat species and in some cases, conducive to human SARS-CoV-2 infection (127). ACE2 also appears to be conserved across mammal species, suggesting that the virus may have the potential to establish in a broad range of other mammalian host species (128–130). Furthermore, there is a possibility that human SARS-CoV-2 could infect initially unsuitable host species through mutation events (131).

To mitigate the risk that SARS-CoV-2 establishes novel enzootic reservoirs and becomes endemic outside of China, it is critical to identify susceptible wildlife species and populations and implement policies that limit their exposure to the virus. The US government has, at present, suspended all bat research to prevent humans from infecting and seeding an enzootic reservoir of SARS-CoV-2 in North American bats (132). However, bat research is still active in many other parts of the world, as are other forms of bat-human contact, though many countries have banned bat bushmeat in response to the pandemic (133). Additional policies may be needed to minimize human contact with other potentially susceptible wildlife populations—species with ACE2 receptors predicted to associate with the SARS-CoV-2 spike receptor binding site (RBD), and populations that occupy anthropogenic landscapes (particularly in regions experiencing a high burden of human transmission) and characterized by demographic rates and structuring predicted to facilitate sustained epidemics and enzootic persistence. It is also critical to monitor via surveillance whether these potentially susceptible animals become exposed and begin to demonstrate capacity for between-host transmission. In particular, researchers should monitor domestic animals, which can bridge transmission between human and wildlife populations (45). Notably, efficient SARS-CoV-2 replication and confirmed cases have been reported in cats (129,134) and ACE2 receptors of domestic cattle have been predicted to associate with the SARS-CoV-2 RBD (135).

The risk of enzootic persistence depends on a multitude of factors, many of which are unknown or poorly characterized, such as infectivity, number of spillback events, and the transmission conditions needed for persistence. Here we provide a conceptual framework of ecological factors to begin addressing the challenge of predicting this risk. Considering ecological mechanisms is a first step towards developing targeted intervention strategies. We additionally argue for the utility of modeling in detangling ecological complexity, providing a simulation model of arboviral transmission in New World primates and mosquitoes as an example. Our results indicate that although long extrinsic incubation periods (EIPs) can reduce the probability of enzootic persistence, the long mosquito lifespans that are characteristic of tropical New World sylvatic species may negate this effect—suggesting that differences in EIP that we may have expected to be important in determining the translocated arboviruses’ relative risk of enzootic establishment are unlikely to be relevant. Overall, our work is important in highlighting the need to be vigilant of imported zoonoses and emphasizing the importance of robust programs to mitigate the risk of spillback events that lead to enzootic persistence.

### Learning points

- The globalization of zoonoses—capable of both infecting humans and establishing persistent enzootic reservoirs of transmission in new regions—poses a unique and critical threat to human health. Understanding the risk that translocated zoonoses spill back and establish reservoirs in novel wildlife hosts should be established as a research priority, particularly in the era of SARS-CoV-2.
- Novel enzootic cycles occur when zoonoses are introduced to new regions (translocation), infect local wildlife (spillback), and persist in local animal host populations (enzootic establishment).
- Understanding when and why translocated zoonoses establish novel enzootic cycles requires disentangling ecologically complex and stochastic interactions between the zoonosis, the human population, and the natural ecosystem.
- Mathematical modeling can inform risk assessments by leveraging existing empirical data. For example, simulation modeling indicates that long extrinsic incubation periods (EIPs) in the mosquito can reduce the probability of enzootic persistence, but the long mosquito lifespans that are characteristic of tropical New World sylvatic species may negate this effect. These model predictions suggest that while we may have expected differences in EIP to significantly affect the risk that translocated arboviruses establish novel enzootic cycles in the neotropics, these differences are unlikely to be relevant.

### Key papers in the field

- Althouse BM, Vasilakis N, Sall AA, Diallo M, Weaver SC, Hanley KA. Potential for Zika Virus to Establish a Sylvatic Transmission Cycle in the Americas. PLOS Neglected Tropical Diseases. 2016 Dec 15;10(12):e0005055.
- Weaver SC. Urbanization and geographic expansion of zoonotic arboviral diseases: mechanisms and potential strategies for prevention. Trends in Microbiology. 2013 Aug;21(8):360-3.
- Han BA, Majumdar S, Calmon FP, Glicksberg BS, Horesh R, Kumar A, et al. Confronting data sparsity to identify potential sources of Zika virus spillover infection among primates. Epidemics. 2019 Jun;27:59-65.
- Lourenço-de-Oliveira R, Failloux A-B. High risk for chikungunya virus to initiate an enzootic sylvatic cycle in the tropical Americas. Vasilakis N, editor. PLOS Neglected Tropical Diseases. 2017 Jun 29;11(6):e0005698.
- Hanley KA, Monath TP, Weaver SC, Rossi SL, Richman RL, Vasilakis N. Fever versus fever: The role of host and vector susceptibility and interspecific competition in shaping the current and future distributions of the sylvatic cycles of dengue virus and yellow fever virus. Infection, Genetics and Evolution. 2013 Oct;19:292-311.

## Supporting Information Legends

### Supplementary Information

An extended description of the simulation model discussed in this manuscript.

## Supporting Information

### Simulation model methods

Our approach builds on Althouse et al. (1), which explored how the force of infection and primate birthrate affect the probability that ZIKV will establish a sylvatic cycle in the Americas. Expanding their analysis, we explored the effects of the mosquito extrinsic incubation period (EIP) and lifespan, and primate birthrate, and considered DENV, CHIKV, and YFV in addition to ZIKV. We simulate the introduction of a single infected primate into metapopulations of susceptible primate hosts and mosquito vectors, applying the Gillespie stochastic simulation algorithm with the Binomial Tau Leap approximation to the following transition rates for mosquitoes:

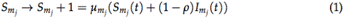

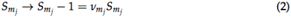

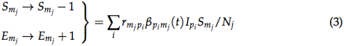

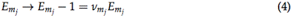

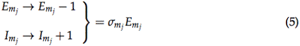

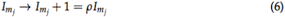

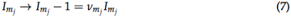

and for primates:

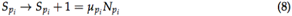

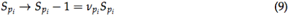

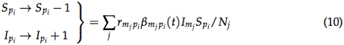

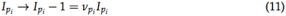

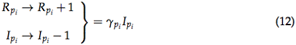

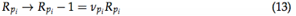

The equations divide primate and mosquito metapopulations into primate species 1,…i and mosquito species 1,…j. The model assumes host preference, with mosquito species j biting the corresponding primate species j more often than other primate species—characterized by the on-diagonal rates within a contact matrix. Primate-mosquito species pairs are coupled by off-diagonal cross-biting rates, calculated as a fixed fraction (10%) of the on-diagonal within-pair biting rates. Given that vertical transmission in the mosquito population is thought to be negligible in enzootic cycles (2), the mosquito transovarial transmission rate *ρ* is set to zero. Thus, all primates and mosquitoes are born susceptible to arbovirus infection at rates *μ* and infected at rates *β*(t), proportional to the number of mosquito bites given or received per day and the probability of successful transmission. We assume that birthrate = 1/lifespan—a conservative estimate given that primates’ peak reproductive years occur before the age of mortality (3) and mosquitoes have the capacity to complete multiple gonotrophic cycles within a lifetime (4). Thus, birthrates vary as we explore three mosquito lifespans (7, 14, and 21 days). Transmission probabilities vary seasonally in Equation 14 and 15 due to changes in environmental conditions such as rainfall and temperature (5). After infection, mosquitoes enter the exposed compartment, progressing to the infectious period at one of three fixed rates *σ_m_j__*. (1/2, 1/7, or 1/10 days).

Primates become infectious immediately and recover at a fixed rate *γ_p_i__*., whereas mosquitoes remain infectious for life. Within each simulation, all parameter values are held constant across mosquito and primate species. Between simulations, only the EIP, and mosquito and primate lifespans, and number of species are varied—all other parameter values are fixed. Holding parameters constant whenever possible allowed us to minimize model complexity and target the effects of our parameters of interest. Furthermore, for the majority of these parameters, there is not enough empirical data to accurately characterize differences between New World species. Full lists of parameter values used for the simulations presented in Figure 2 are given in the table below. For each set of parameter values, we ran 50 simulations and calculated the probability of sylvatic establishment as the proportion of simulations in which there were infected primates remaining at the end of the simulated three-year period.

| Parameter | Description | Value |
| --- | --- | --- |
| $t$ | Time in days | |
| $r_{m_j p_i}$ | Biting rate per day to primates by mosquito $j$ | $0.5 \text{ days}^{-1}$ |
| $b_{p_i m_j}$ | Baseline transmission probability, from primate $i$ to mosquito $j$ | 0.6 |
| $b_{m_j p_i}$ | Baseline transmission probability, from mosquito $j$ to primate $i$ | 0.6 |
| $c_j$ | Percent of the magnitude of seasonal variation for mosquito $j$ | 0.1 |
| $\mu_{p_i}$ | Primate $i$ birth rate (=1/lifespan) | $1/(5 \times 365)$ to $1/(15 \times 365) \text{ years}^{-1}$ |
| $\nu_{p_i}$ | Primate $i$ death rate, set equal to birth rate | |
| $\gamma_{p_i}$ | Primate $i$ recovery rate | $1/4 \text{ days}^{-1}$ |
| $\mu_{m_j}$ | Mosquito $j$ birth rate | $1/7 \text{ days}^{-1}$ |
| $\nu_{m_j}$ | Mosquito $j$ death rate, set equal to birth rate | |
| $\sigma_{m_j}$ | Extrinsic incubation period in mosquito $j$ | 0 |
| $\rho$ | Mosquito rate of transovarial transmission | 0 |

By default, the compartmental model described in Equation 1 through 13 assumes exponentially distributed waiting periods. This assumption is biologically unrealistic because individuals leave their current compartment at a constant rate, regardless of how much time has passed since they entered that compartment. For example, with constant rates, individuals have an equal probability of leaving the latent compartment regardless of the time since infection. In reality, the probability of progressing should be very low immediately after infection, and highest around the latent period mean—a trajectory better represented by a gamma distribution, or it’s discrete analog, the Erlang distribution. The use of exponentially distributed infectious and latent periods significantly affects model estimations of the basic reproductive number, duration of epidemics, and critical population size required for disease persistence in directly transmitted pathogens (6–8). Given the short mosquito lifespan and the resulting interaction between EIP and the length of the infectious period, epidemic dynamics would be more sensitive to the distribution of waiting periods in the mosquito vector than in the primate host. Thus, we explored the effect of modeling the latent period in the mosquito vector (EIP) as an Erlang distribution by splitting the latent period into a series of separate compartments, commonly referred to as a boxcar configuration (9).

We constructed two models with Erlang-distributed EIPs—a model with 10 exposed compartments, and a model with 50 exposed compartments. Figure S1 demonstrates the differences between the probability distributions of the exponentially-distributed EIP relative to both Erlang-distributed EIPs. As expected, modeling EIP as less-dispersed Erlang distributions resulted in higher probabilities of longer EIPs, thus decreasing the probability of sylvatic persistence. However, the Erlang-distributed models produced the same general trends as the exponentially distributed models, with shorter EIPs and longer mosquito lifespans increasing the probability of sylvatic establishment. As a result, in the interest of minimizing model complexity, in our main text, we report results from our exponentially-distributed model. Simulation results from the Erlang-distributed models with respect to differences in the mosquito EIP and lifespan are presented in Figure S2 and S3.

**Figure S1.**
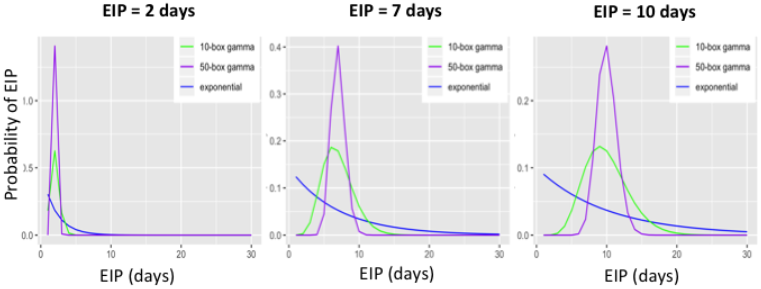
Comparison of exponential- and gamma-distributed EIPs. By default, waiting periods (such as the EIP) in traditional compartment models follow exponential probability distributions (in blue). In the case of EIP, this assumption is unrealistic because individuals have an equal probability of leaving the incubation compartment regardless of the time since infection. In reality, the probability of progressing should be very low immediately after infection and highest around the EIP mean—a trajectory better represented by a gamma distribution (in green and purple). A gamma distributed EIP can be constructed in a compartmental model via a boxcar configuration of the latent period (i.e., splitting the latent period up into a series of separate compartments, or boxes). Note how as the number of latent boxes increases (green vs. purple), the gamma distribution becomes less dispersed, and more closely centered around the EIP mean.

**Figure S2.**
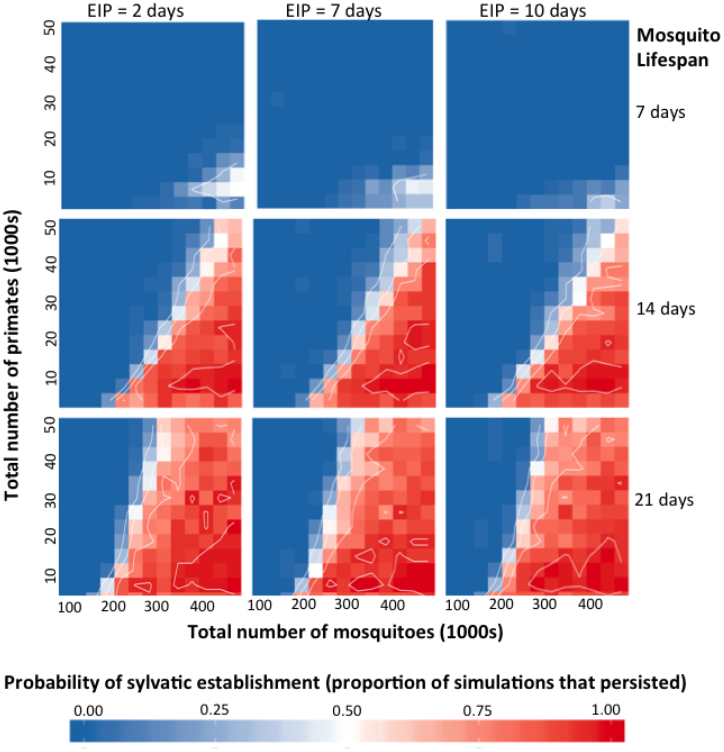
The probability of establishment with a 10-box Erlang-distributed EIP. Mosquito birthrate = 1/lifespan and increases from the top to bottom panels, while EIP increases left to right. Here, we constructed an Erlang-distributed EIP by splitting the exposed compartment into 10 separate boxes. Within each panel, the total population size of mosquitoes (in two populations) and primates (in two populations) changes horizontally and vertically, respectively. For each parameter set, we simulated the introduction of a single infected primate and subsequent transmission for a three-year period. Blue indicates no simulations establishing, whereas red indicates all simulations establishing. Contour lines show 0.25, 0.5, 0.75, and 0.95 probability of establishment.

**Figure S3.**
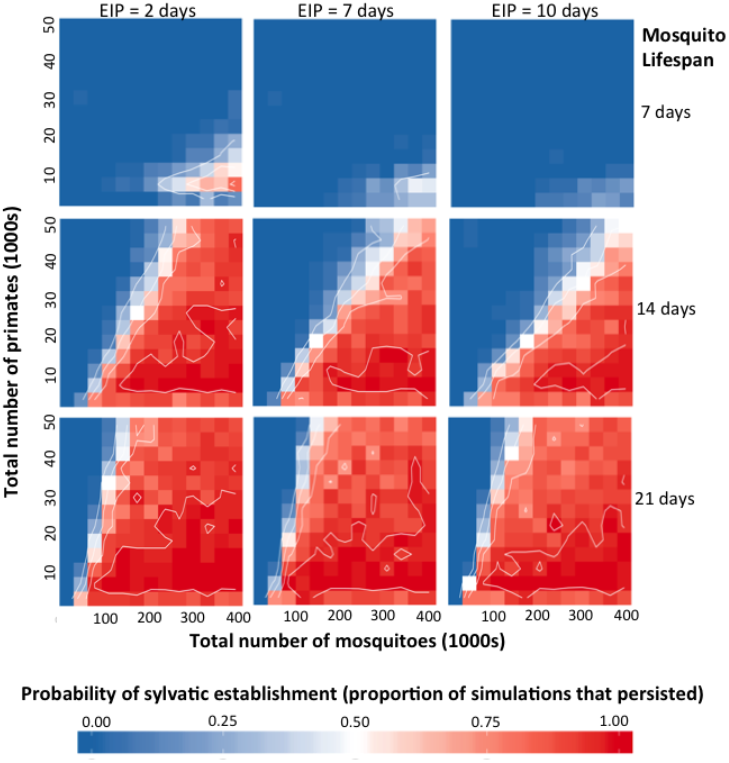
The probability of establishment with a 50-box Erlang-distributed EIP. Mosquito birthrate = 1/lifespan and increases from the top to bottom panels, while EIP increases left to right. Here, we constructed an Erlang-distributed EIP by splitting the exposed compartment into 50 separate boxes. Within each panel, the total population size of mosquitoes (in two populations) and primates (in two populations) changes horizontally and vertically, respectively. For each parameter set, we simulated the introduction of a single infected primate and subsequent transmission for a three-year period. Blue indicates no simulations establishing, whereas red indicates all simulations establishing. Contour lines show 0.25, 0.5, 0.75, and 0.95 probability of establishment.

## Notes

### Competing Interest Statement

The authors have declared no competing interest.

